# AP-4 mediates vesicular transport of the 2-AG endocannabinoid producing enzyme DAGLB

**DOI:** 10.1101/2020.10.25.353995

**Authors:** Alexandra K. Davies, Marvin Ziegler, Hellen Jumo, Wardiya Afshar Saber, Darius Ebrahimi-Fakhari, Georg H. H. Borner

## Abstract

The adaptor protein complex AP-4 mediates anterograde axonal transport and is essential for axon health. AP-4-deficient patients suffer from a severe neurological disorder, which encompasses neurodevelopmental and neurodegenerative features. While impaired autophagy has been suggested to account for axon degeneration in AP-4 deficiency, axon growth defects occur through an unknown mechanism. Here we use orthogonal proteomic and imaging methods to identify DAGLB (diacylglycerol lipase-beta) as a cargo of AP-4 vesicles. DAGLB is a key enzyme for the generation of 2-AG (2-arachidonoylglycerol), the most abundant endocannabinoid in brain. During normal development, DAGLB is targeted to the axon, where 2-AG signalling drives axonal growth. We show that DAGLB accumulates at the TGN of AP-4-deficient cells, including in iPSC-derived neurons from a patient with AP-4 deficiency syndrome. Our data thus support that AP-4 mediates axonal targeting of DAGLB, and we propose that axon growth defects in AP-4 deficiency may arise through spatial dysregulation of endocannabinoid signalling.

## Introduction

Axon development and maintenance critically rely on tightly regulated anterograde transport. The factors that promote axonal growth and the machinery that removes dysfunctional components must efficiently reach the distal axon. Accordingly, mutations in the axonal transport machinery are implicated in a broad range of neurodevelopmental and neurodegenerative diseases (reviewed in Guedes-Dias & Holzbaur, 2019). We and others recently discovered that adaptor protein complex 4 (AP-4) is required for the axonal delivery of vesicles containing the autophagy protein ATG9A (Davies et al., 2018; De Pace et al., 2018; Ivankovic et al., 2020; Mattera et al., 2017). Biallelic loss-of-function mutations in any of the four subunits of AP-4 (*AP4B1, AP4E1, AP4M1, AP4S1*) cause AP-4 deficiency syndrome, characterised by global developmental delay, intellectual disability, seizures and progressive spasticity, with onset of symptoms in early infancy (Abou Jamra et al., 2011; Moreno-De-Luca et al., 2011; Verkerk et al., 2009). There is currently no treatment, and a steep rise in diagnoses over the last three years (Ebrahimi-fakhari et al., 2020) is fuelling efforts to understand the mechanisms that cause neuronal pathology in AP-4 deficiency syndrome.

Neuron-specific knockout of the core autophagy genes *Atg5* or *Atg7* in mice leads to neurodegeneration (Hara et al., 2006; Komatsu et al., 2006). Therefore, the current model for AP-4 deficiency suggests that lack of ATG9A in the distal axon leads to impaired autophagosome biogenesis, causing axonal degeneration. However, defects in neuronal autophagy are insufficient to explain all aspects of the disease. *In vitro*, neurons from embryonic *Ap4e1* knockout mice have axonal growth defects including reduced length and branching (Ivankovic et al., 2020). In contrast, axon length is normal in cultured neurons from autophagy-deficient *Atg7* knockout mouse embryos (Yamaguchi et al., 2018). This suggests that AP-4 has an unidentified role in neurodevelopment that is independent of its role in autophagy.

Unbiased proteomic approaches have yielded a consensus set of AP-4 vesicle cargoes and accessory proteins (summarised in Table S1; Borner et al., 2012; Davies et al., 2018; Mattera et al., 2015). Our spatial proteomics method, Dynamic Organellar Maps, provided a powerful hypothesis-free screen for proteins with an altered distribution in AP-4-deficient cells, leading to the identification of the AP-4 cargo proteins ATG9A, SERINC1 and SERINC3 (Davies et al., 2018). Using this approach, we here identify DAGLB (diacylglycerol lipase-beta) as a cargo protein of AP-4 vesicles. DAGLB is a key enzyme for generation of 2-arachidonoyl-glycerol (2-AG), the most abundant endocannabinoid in brain. In developing neurons, DAGLB is targeted to the distal axon by an unknown mechanism, where 2-AG is required to promote axonal growth and guidance (Bisogno et al., 2003; Williams et al., 2003). Our data support that AP-4 mediates anterograde axonal transport of DAGLB. We therefore suggest that AP-4 provides a missing link in the regulation of endocannabinoid signalling, and that neurodevelopmental aspects of AP-4 deficiency syndrome may arise through disruption of the spatial control of 2-AG synthesis.

## Results and discussion

### Sensitive spatial proteomics reveals DAGLB as an AP-4 cargo protein

We previously used our spatial proteomics method, Dynamic Organellar Maps, to identify cargoes of the AP-4 vesicle pathway (Davies et al., 2018). In this approach, a ‘map’ of the cell is generated by combining cell fractionation with MS-based quantification to provide proteome-wide protein localisation information (Itzhak et al., 2016). We compared maps from wild-type, *AP4B1* knockout and *AP4E1* knockout HeLa cells in biological duplicate, to look for proteins with an altered subcellular localisation in the AP-4-deficient cells. Using very stringent data filters (false discovery rate (FDR) < 1%), we identified three proteins with a localisation shift – ATG9A, SERINC1 and SERINC3 – which we then validated as *bona fide* AP-4 cargoes by imaging (Davies et al., 2018).

To investigate if AP-4 may have additional cargo proteins, we now performed a more sensitive exploratory analysis of our data (Fig. 1a; see methods for details). We identified eight additional hits, with an estimated FDR of approximately 25% (inset plot of Fig. 1a). As a further filter we visualised the localisation of our new hits on an organellar map of wild-type HeLa cells (Fig. 1b; Itzhak et al., 2016). Three proteins, DAGLB, PTPN9 and LNPEP, mapped to the endosomal cluster, like the known AP-4 cargo proteins. Therefore, they are candidate novel AP-4 vesicle proteins (Table S1). Of note, DAGLB and PTPN9 were found in close proximity to ATG9A.

**Figure 1.**
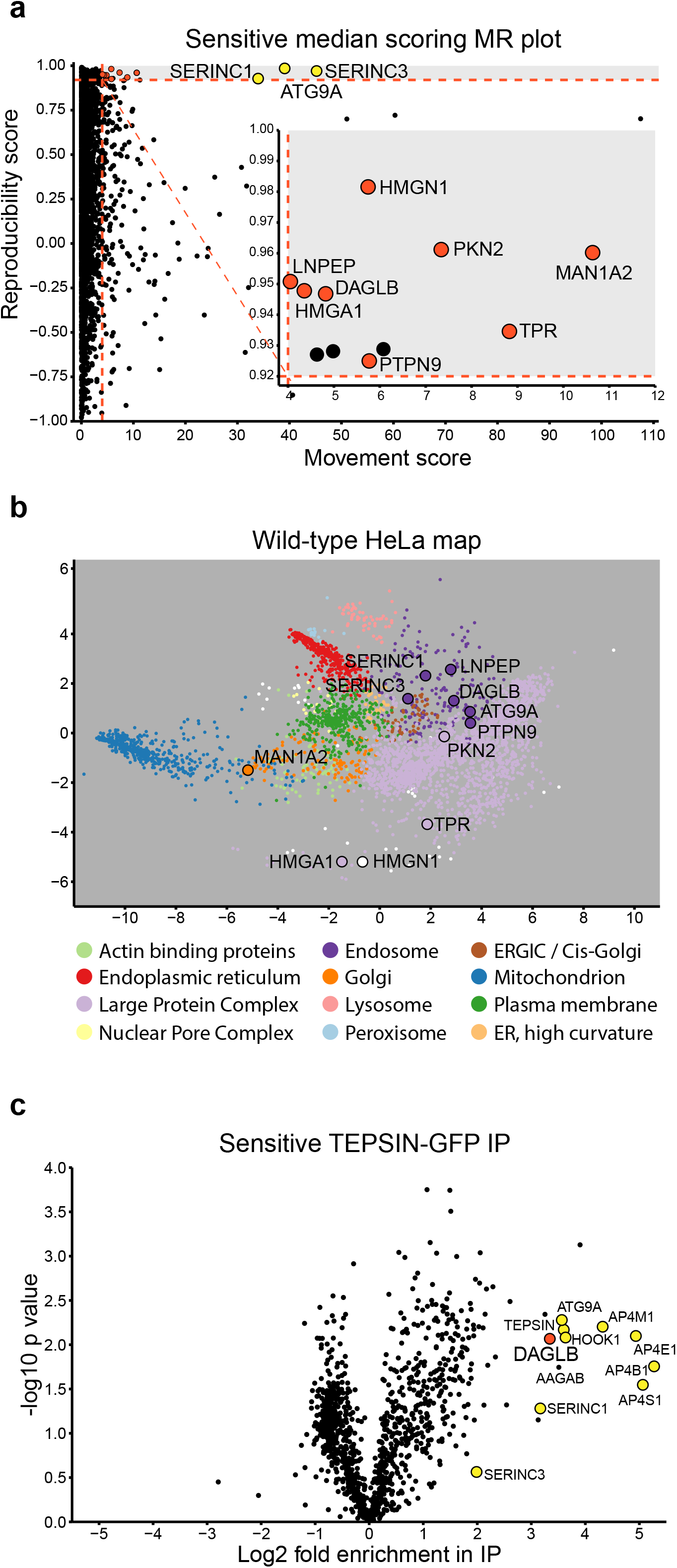
Sensitive analysis of Dynamic Organellar Maps identifies DAGLB as an AP-4 cargo protein. **(a)** Dynamic Organellar Maps of *AP4B1* knockout (KO) and *AP4E1* KO HeLa cells were compared to maps of wild-type HeLa cells (each in duplicate, totalling 4 comparisons). Sensitive statistical scoring was used to detect proteins with a significantly altered distribution in the AP-4 KO cells that were not detected in a previous stringent analysis (see methods section for details; Davies et al., 2018). For 3926 proteins profiled across all maps, the ‘MR’ plot displays the median magnitude of shift (M) and the mean within-clone reproducibility of shift direction (R). The known AP-4 cargo proteins, ATG9A, SERINC1 and SERINC3, were identified with high M and R scores, as expected. The inset plot displays 8 additional hits (marked in red) whose subcellular localisation was significantly and reproducibly shifted in the AP-4 KO lines, with a false discovery rate of ~25%. **(b)** The hits from **a** are highlighted on a principal component analysis (PCA)-based visualisation of a deep Dynamic Organellar Map of wild-type HeLa cells (data from 6 maps; Itzhak et al., 2016). Each scatter point represents a protein and proximity indicates similar fractionation profiles. Colours indicate subcellular compartment assignment by support vector machine-based classification (white indicates compartment unassigned). Three hits, DAGLB, PTPN9 and LNPEP, map to the endosomal cluster (dark purple), like known AP-4 cargo proteins, making them strong candidates for AP-4 vesicle proteins. **(c)** High-sensitivity low-detergent immunoprecipitations (IP) from HeLa cells stably expressing the AP-4 associated protein TEPSIN-GFP were analysed by SILAC-based quantitative mass spectrometry (Davies et al., 2018). Data were analysed in comparison to mock immunoprecipitations from wild-type HeLa cells with a two-tailed one sample *t*-test (each in triplicate, *n* = 3). DAGLB (marked in red) was highly enriched along with known AP-4 vesicle proteins (marked in yellow).

In our previous study, we found that ATG9A, SERINC1 and SERINC3 were co-immunoprecipitated with the AP-4 complex under sensitive low-detergent conditions (Davies et al., 2018). We re-analysed this proteomic dataset and found that DAGLB was indeed co-enriched, along with known AP-4 vesicle proteins (Fig. 1c). Combined with our mapping data, this strongly supports that DAGLB is an AP-4 cargo protein. DAGLB is a serine lipase that hydrolyses diacylglycerol (DAG) to generate the signalling lipid 2-AG (Bisogno et al., 2003). 2-AG is an agonist for the CB1 and CB2 cannabinoid receptors, which have important functions in axonal growth and guidance during development (reviewed in Oudin et al., 2011). As axonal growth defects arise through an unknown mechanism in AP-4-deficient neurons, we investigated AP-4-dependent localisation of DAGLB in detail.

### DAGLB accumulates at the TGN of AP-4 deficient HeLa and neuronal SH-SY5Y cells

In the absence of AP-4, ATG9A accumulates at the TGN of diverse cell types, including neuronal and nonneuronal cells (Behne et al., 2020; Davies et al., 2018; De Pace et al., 2018; Ivankovic et al., 2020; Mattera et al., 2017). To assess whether DAGLB exhibits a similar missorting phenotype in AP-4-deficient cells, we used immunofluorescence microscopy. In wild-type cells DAGLB was observed in fine puncta throughout the cell, with increased density in the region of the TGN (Fig. 2a), a strikingly similar distribution to that of ATG9A (Davies et al., 2018; Young et al., 2006). In *AP4B1* knockout cells there was a significant increase in the ratio of DAGLB intensity between the TGN and the rest of the cell, and this was rescued by stable expression of the missing AP4B1 subunit (Fig 2a and b). A similar increase in the amount of DAGLB at the TGN was observed in cells treated with siRNA to knock down AP-4 (Fig. S1a and b). These data suggest that AP-4 deficiency causes a partial block of DAGLB export from the TGN. The effect was not as striking as that observed for ATG9A (Davies et al., 2018), but this is consistent with the lower movement score of DAGLB in our Dynamic Organellar Maps analysis, and highlights the sensitivity of our proteomic method for detecting even very subtle protein localisation changes. Next, we examined DAGLB distribution in neuroblastoma-derived SH-SY5Y cells depleted of AP4B1 or AP4E1 using CRISPR/Cas9 (Davies et al., 2018), differentiated into a neuron-like state by the addition of retinoic acid and BDNF. In the AP-4 depleted SH-SY5Y cells there was also an increase in the ratio of DAGLB at the TGN compared to control cells (Fig. 2c and d). This was more pronounced than in HeLa cells, indicating that AP-4 dependent sorting of DAGLB may be particularly important in polarised cells. The ratio of DAGLB at the TGN was also increased in AP-4-depleted SH-SY5Y cells in their undifferentiated state (Fig. S1c and d). Collectively, these observations confirm DAGLB as an AP-4 cargo in HeLa and SH-SY5Y cells, and suggest that TGN export of DAGLB is a ubiquitous function of the AP-4 pathway.

**Figure 2.**
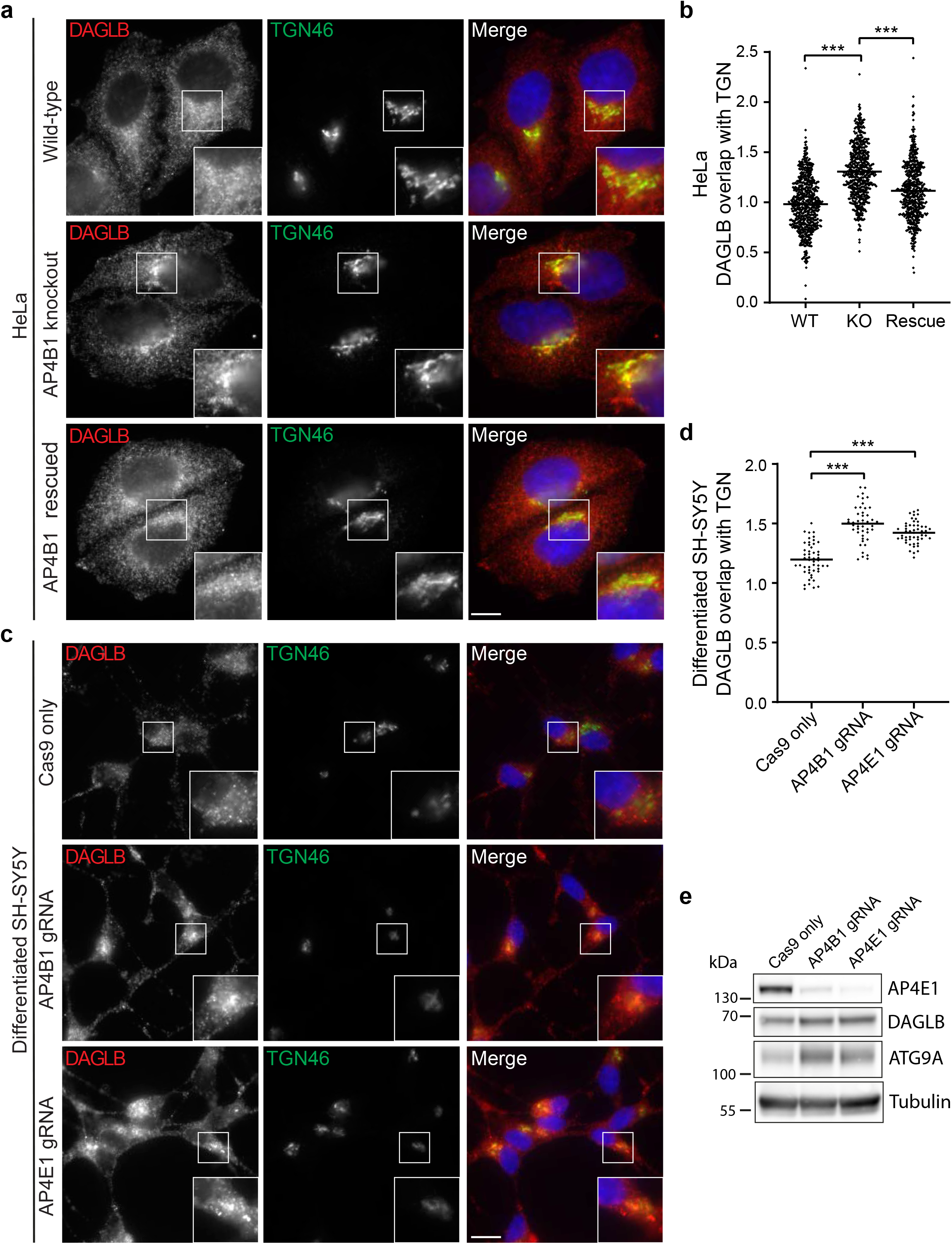
DAGLB accumulates at the *trans*-Golgi network (TGN) in AP-4 knockout HeLa and differentiated SH-SY5Y cells. **(a)** Widefield imaging of immunofluorescence double labelling of DAGLB and TGN46 in wild-type, *AP4B1* knockout, and *AP4B1* knockout HeLa cells stably expressing AP4B1 (functional rescue). In the merged image, DAPI labelling of the nucleus is also shown (blue). Scale bar: 10 μm. **(b)** Quantification of the ratio of DAGLB labelling intensity between the TGN and the rest of the cell, in the cells shown in **a**. The experiment was performed in biological triplicate, and >150 cells were scored per cell line in each replicate. The graph shows combined replicate data, and each datapoint indicates the ratio for an individual cell (horizontal bar indicates mean). Data were subjected to a Kruskal-Wallis test with Dunn’s Multiple Comparison Post-Test for significance: *\*\*\*p* ≤ 0.001. **(c)** CRISPR-Cas9 was used to deplete AP4B1 or AP4E1 in mixed populations of SH-SY5Y cells (Davies et al., 2018). Widefield imaging of immunofluorescence double labelling of DAGLB and TGN46 in control (parental Cas9-expressing), AP4B1-depleted and AP4E1-depleted neuronally-differentiated SH-SY5Y cells. In the merged image, DAPI labelling of the nucleus is also shown (blue). Scale bar: 10 μm. **(d)** Quantification of the ratio of DAGLB labelling intensity between the TGN and the rest of the cell, in the cells shown in **c**. The experiment was performed in biological triplicate, and >10 images were quantified per cell line in each replicate. The graph shows combined replicate data, and each datapoint indicates the ratio calculated from a single image (horizontal bar indicates mean). Data were subjected to a Kruskal-Wallis test with Dunn’s Multiple Comparison Test for significance: \*\*\**p* ≤ 0.001. **(e)** Western blots of whole cell lysates from the cells shown in **c** & **d**; Tubulin, loading control. Representative of two independent experiments.

AP-4-deficiency increases the expression level of ATG9A in neurons (De Pace et al., 2018; Ivankovic et al., 2020) and patient fibroblasts, and this serves a robust diagnostic disease marker (Behne et al., 2020). Likewise, we observed increased levels of DAGLB in whole cell lysates from AP-4-depleted neuronally differentiated SH-SY5Y cells (Fig. 2e). As has been suggested for ATG9A, this could represent an adaptive mechanism to compensate for the lack of directional transport of DAGLB. Alternatively, it is possible that the altered trafficking itinerary of ATG9A and DAGLB results in reduced turnover of the proteins. Mattera and colleagues (2017) did not find any difference in the stability of ATG9A over 20 hours in a cycloheximide chase experiment, but a longer time course may be necessary. Regardless, the fact that DAGLB exhibits behaviour similar to ATG9A in response to lack of AP-4 further supports that it is an AP-4 cargo protein.

### DAGLB colocalizes with AP-4 vesicle proteins

The AP-4 accessory protein RUSC2 mediates microtubule plus-end-directed transport of AP-4 vesicles from the TGN, presumably by recruiting a kinesin molecular motor (Davies et al., 2018). Overexpression of RUSC2 dramatically alters the localisation of the AP-4 cargo proteins ATG9A, SERINC1 and SERINC3 to the periphery of the cell, and this only occurs in cells that express AP-4. We exploited this behaviour to test if DAGLB is present in AP-4-derived vesicles. Supporting our hypothesis, DAGLB colocalised with RUSC2 in bright puncta at the periphery of HeLa cells that stably overexpress RUSC2-GFP (Fig. 3). Importantly, this was completely dependent on AP-4 – in *AP4B1* knockout cells neither DAGLB nor RUSC2 accumulated at the cell periphery, and they also no longer colocalised with each other. Transient expression of AP4B1 in the *AP4B1* knockout rescued both the peripheral localisation and the colocalisation between DAGLB and RUSC2. This demonstrates that DAGLB is present in AP-4 vesicles, which bud from the TGN and also contain ATG9A, SERINC1 and SERINC3 (Davies et al., 2018).

**Figure 3.**
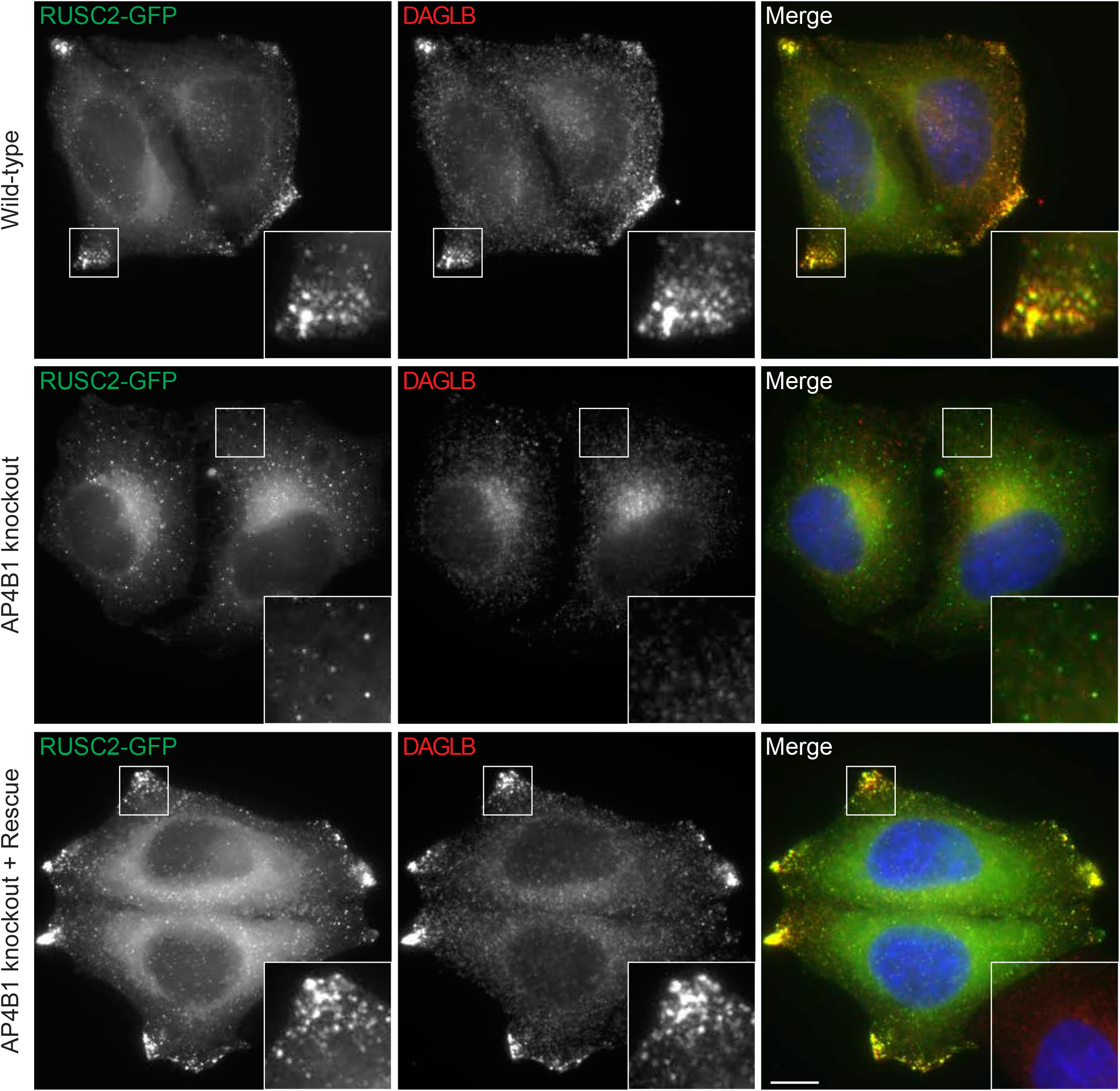
Overexpression of the AP-4 vesicle transport adaptor RUSC2 drives DAGLB to the cell periphery. Widefield imaging of HeLa cells stably expressing RUSC2-GFP, labelled with anti-DAGLB (red channel). Top panel, wild-type; middle panel, *AP4B1* knockout; lower panel, *AP4B1* knockout with transient expression of AP4B1 (rescue). In the merged image, DAPI labelling of the nucleus is also shown (blue). The insets show accumulation of RUSC2-GFP-positive and DAGLB-positive puncta at the cell periphery, and this only occurs in the presence of AP-4 (wild-type and rescue). Scale bar: 10 μm.

To further confirm the presence of DAGLB in AP-4 vesicles, we used super-resolution structured illumination microscopy (SR-SIM) to visualise colocalisation between AP-4 vesicle cargoes in HeLa cells. To calibrate our imaging pipeline, we first quantified colocalisation between the known AP-4 cargo proteins SERINC1 and ATG9A. As expected, there was considerable overlap between SERINC1 and ATG9A in small peripheral puncta, and colocalisation was significantly reduced by AP-4 knockdown (Fig. 4a and b), consistent with our previous analysis (Davies et al., 2018). DAGLB and ATG9A could not be assessed in the same cells due to antibody incompatibility, so we analysed colocalisation between DAGLB and SERINC1 (Fig. 4c and d). The results were extremely similar to those observed for ATG9A. DAGLB colocalised with SERINC1 in small peripheral puncta, and AP-4 knockdown resulted in loss of these puncta. To summarise, our RUSC2 overexpression and SIM colocalisation assays demonstrate that DAGLB is a cargo in AP-4 vesicles.

**Figure 4.**
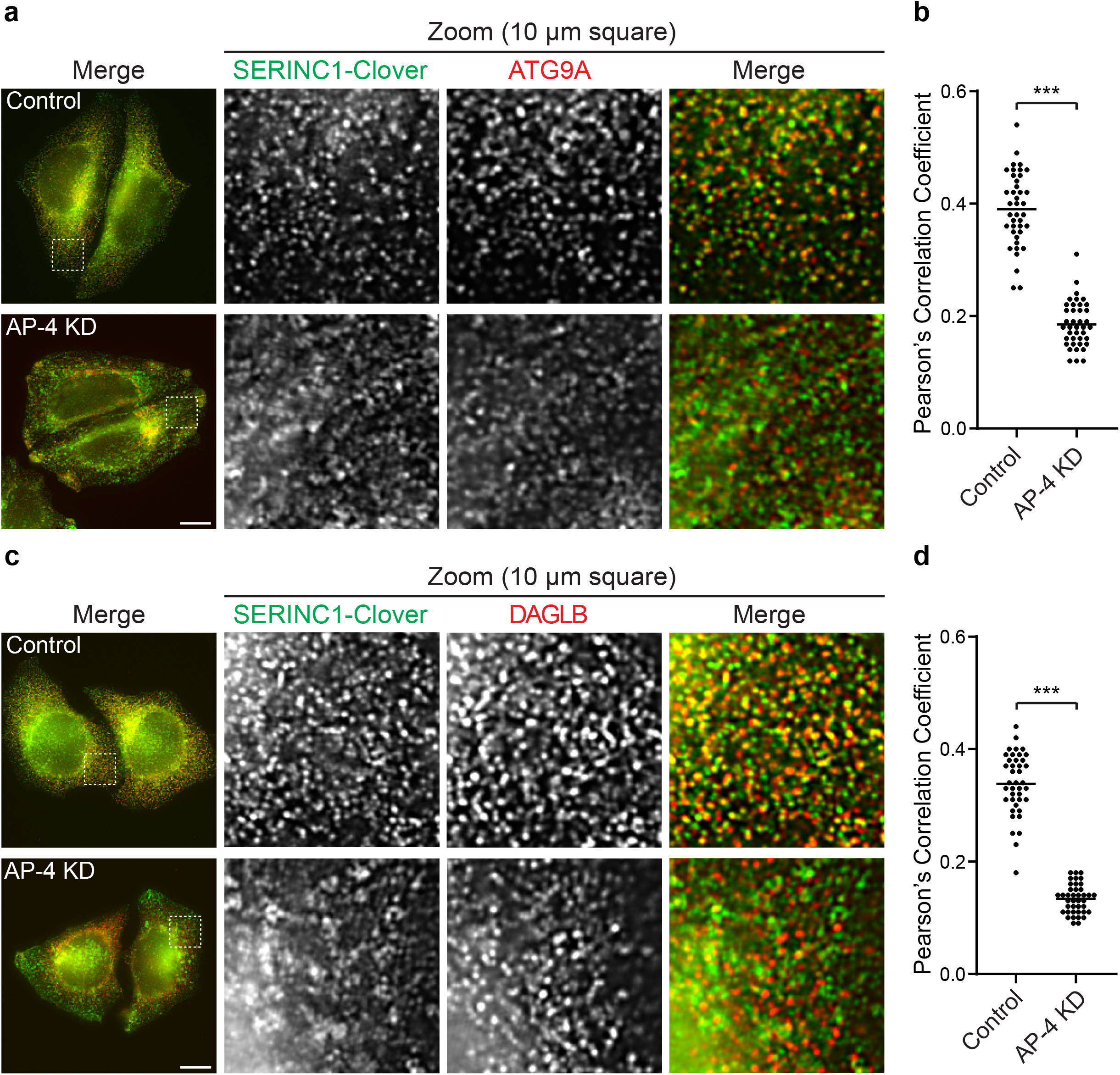
DAGLB and ATG9A colocalise with SERINC1 in an AP-4-dependent manner. HeLa cells tagged endogenously with Clover (modified GFP) at the C-terminus of SERINC1 were transfected with siRNA to knock down (KD) AP-4, or were transfected with a non-targeting siRNA (Control). **(a)** Super resolution structured illumination microscopy (SR-SIM) was used to image SERINC1-Clover (via anti-GFP) and anti-ATG9A. Representative images show the whole field of view and a zoomed image of a peripheral 10 × 10 μm square. ATG9A and SERINC1 colocalise in small puncta throughout the cytoplasm in control cells, but not in AP-4 depleted cells. Scale bar: 10 μm. **(b)** Quantification of colocalisation between SERINC1-Clover and ATG9A in control and AP-4 knockdown (KD) cells, using Pearson’s Correlation Coefficient (PCC). The experiment was performed in biological duplicate, and 20 cells were scored per condition in each replicate. The graph shows combined replicate data, and each datapoint indicates the PCC for an individual cell (horizontal bar indicates mean). Data were subjected to a two-tailed Mann–Whitney *U*-test: \*\*\**p* ≤ 0.001. **(c)** SR-SIM was used to image SERINC1-Clover (via anti-GFP) and anti-DAGLB, as in **a**. Like ATG9A, DAGLB colocalises with SERINC1 in small puncta throughout the cytoplasm in control cells, but not in AP-4 depleted cells. **(d)** Quantification of colocalisation between SERINC1-Clover and DAGLB in control and AP-4 KD cells, using PCC. The experiment was performed in biological duplicate, and 20 cells were scored per condition in each replicate. The graph shows combined replicate data, and each datapoint indicates the PCC for an individual cell (horizontal bar indicates mean). Data were subjected to a two-tailed Mann–Whitney *U*-test: \*\*\**p* ≤ 0.001.

### DAGLB is mislocalised in iPSC-derived neurons from AP-4-deficient patients

We recently developed the first human neuron model of AP-4 deficiency syndrome, using iPSCs from patients with AP4B1 deficiency (SPG47), differentiated into excitatory cortical neurons (Behne et al., 2020). As in primary neurons from *Ap4e1* knockout mice (De Pace et al., 2018; Ivankovic et al., 2020), ATG9A accumulates at the TGN of the AP-4-deficient human neurons, and is elevated at the whole cell level (Behne et al., 2020). To test if DAGLB shows similar alterations, we compared the distribution of DAGLB relative to TGN46 in iPSC-derived neurons from a patient with AP4B1 deficiency/SPG47 and their same-sex parent, an unaffected heterozygous carrier (Fig. 5a and b). Indeed, DAGLB accumulated at the TGN of neurons from the AP-4-deficient patient, in a similar manner to ATG9A. This provides strong evidence that DAGLB missorting occurs in neurons of patients with AP-4 deficiency syndrome.

**Figure 5.**
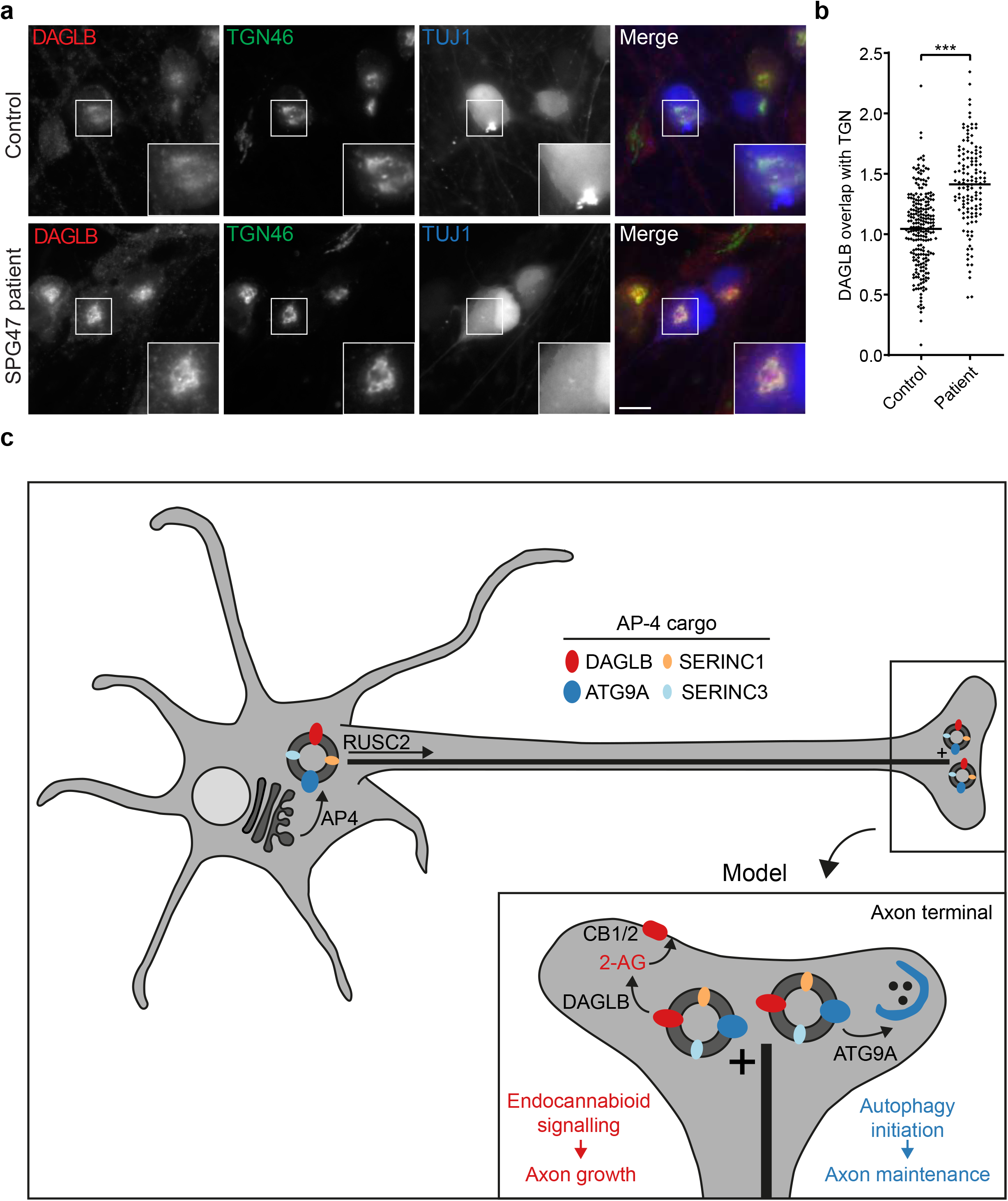
DAGLB is missorted in iPSC-derived cortical neurons from an AP-4 patient, suggesting a new model for AP-4 deficiency syndrome. **(a)** Induced pluripotent stem cells (iPSCs) from a patient with AP4B1-associated AP-4 deficiency syndrome (SPG47; *AP4B1*, NM_001253852.1: c.1345A>T / c.1160_1161delCA) and their unaffected same sex heterozygous parent were differentiated into cortical neurons. Widefield imaging of immunofluorescence triple labelling of DAGLB, TGN46 and TUJ1 (a marker to distinguish neurons from cocultured astrocytes). Scale bar: 10 μm. **(b)** Quantification of the ratio of DAGLB labelling intensity between the *trans*-Golgi network (TGN) and the rest of the cell, in the cells shown in **a**. The experiment was performed in technical triplicate, and >30 cells were scored per cell line in each replicate. The graph shows combined replicate data, and each datapoint indicates the ratio for an individual cell (horizontal bar indicates mean). Data were subjected to a two-tailed Mann–Whitney *U*-test: \*\*\**p* ≤ 0.001. **(c)** Proposed model for the role of AP-4 in axonal transport based on our new and published data. AP-4 acts at the TGN membrane to package its cargo proteins, DAGLB, ATG9A, SERINC1 and SERINC3, into transport vesicles. RUSC2 mediates microtubule plus-end-directed transport of AP-4-derived vesicles, delivering cargo to the distal axon. DAGLB is an enzyme responsible for production of the endocannabinoid 2-AG, known to be required for axonal growth via autocrine activation of cannabinoid receptors. The previous model for AP-4 deficiency syndrome has focused on missorting of ATG9A, which is required for autophagy initiation and hence for axonal maintenance. We now propose that the neuronal pathology in AP-4 deficiency arises from the compound effects of DAGLB missorting on axonal development and ATG9A missorting on axonal autophagy.

### A new model for AP-4 deficiency syndrome

DAGLB is a key enzyme for generation of the endocannabinoid 2-AG, and DAGLB missorting in AP-4 deficient neurons is hence likely to contribute to the neurodevelopmental aspects of AP-4 deficiency syndrome. 2-AG is a key signalling lipid in the central nervous system. The major pathway for 2-AG biosynthesis is hydrolysis of DAG at the *sn1* position by one of two closely related DAG lipases – DAGLA and DAGLB (Bisogno et al., 2003). DAGLs are required for diverse aspects of brain function, including axonal growth during development, and neurogenesis and retrograde synaptic signalling in the adult brain (Oudin et al., 2011). Crucially, these diverse functions require not only temporal control of DAGL expression, but also spatial regulation. In developing neurons, the DAGLs are present in growing axons (Bisogno et al., 2003; Mulder et al., 2008), which correlates with a requirement for locally synthesised 2-AG to promote axonal growth and guidance by autocrine activation of CB1 receptors (Williams et al., 2003). However, despite the importance for axon development, the mechanism of axonal targeting of DAGLs is currently unknown.

AP-4 mediates axonal delivery of ATG9A (Ivankovic et al., 2020). Here we show that DAGLB is present in the same AP-4-derived vesicles as ATG9A (Fig. 3 and 4), and that it accumulates at the TGN of AP-4 patient-derived neurons (Fig. 5a and b). Therefore, our data strongly suggest that AP-4 is responsible for the axonal targeting of DAGLB (Fig. 5c). AP-4-deficient neurons exhibit axon-specific defects including reduced length, reduced branching and axonal swellings (Behne et al., 2020; De Pace et al., 2018; Ivankovic et al., 2020). Although autophagy is implicated in neurodegeneration, aberrant autophagy alone is insufficient to explain reduced axonal growth (Yamaguchi et al., 2018). Brain imaging studies of AP-4-deficient patients show features that suggest loss of axons, in particular of the long projecting neurons, including almost universal thinning of the corpus callosum and ventriculomegaly (enlarged ventricles) in the majority of cases (Ebrahimi-fakhari et al., 2020). Endocannabinoid signalling is required for elongation and fasciculation of the long axons of pyramidal cells (a common class of projection neurons in the cerebral cortex; Mulder et al., 2008). Thus, we propose that dysregulation of the spatial control of 2-AG production may contribute to axon defects in AP-4 deficiency syndrome (Fig. 5c).

An important open question is whether AP-4 also transports DAGLA or whether there is differential sorting of the two DAG lipases. Both enzymes are important contributors to 2-AG biosynthesis; *Dagla* and *Daglb* knockout mice both have significantly reduced levels of 2-AG in the brain (around 80% and 50% reductions, respectively; Gao et al., 2010). DAGLA is not detectable in our proteomic analyses of HeLa or neuronally differentiated SH-SY5Y cells; hence, we focussed our attention on DAGLB. RNA-based expression analysis suggests that, like other AP-4 vesicle proteins, DAGLB is ubiquitously expressed in human tissues (Uhlén et al., 2015; Human Protein Atlas available from http://www.proteinatlas.org). In contrast, the expression of DAGLA is elevated in the brain and is lower than that of DAGLB in other tissues. We used mass spectrometry to quantify DAGLA and DAGLB in wild-type and *Ap4e1* knockout mouse brains and found that DAGLB is slightly overexpressed in *Ap4e1* knockout brains (approximately 20% increase), whereas DAGLA is unaffected (our unpublished data). This supports the possibility that AP-4 may play a specific role in the sorting of DAGLB. However, future studies should investigate DAGLA distribution in AP-4 deficient cells to test this hypothesis.

In conclusion, we have identified an important role for AP-4 in transport of DAGLB, and thus provide a missing link in understanding the regulation of endocannabinoid signalling. Furthermore, our findings open up a new direction for the pathomechanisms of AP-4 deficiency syndrome – the possibility that aberrant spatial control of 2-AG synthesis leads to developmental defects in axon formation. We suggest that the diverse neurological symptoms of AP-4-deficient patients may be caused by compound effects of DAGLB missorting on axonal development and ATG9A missorting on axonal maintenance. Furthermore, our study predicts that global or local neuronal 2-AG levels may be affected in AP-4 deficiency, and we are currently developing a lipidomics approach to investigate this. A number of pharmacological tools for intervening in 2-AG-mediated endocannabinoid signalling already exist (reviewed in Baggelaar et al., 2018), and an inhibitor of monoacylglycerol lipase (MAGL), the enzyme responsible for degrading 2-AG (Dinh et al., 2002), is currently in clinical trials for treating pain disorders (Egmond et al., 2021). It will be important to test if increasing 2-AG levels through MAGL inhibition might help to correct the axonal defects observed in AP-4 deficient cells, which may enable new treatment strategies for AP-4 deficiency syndrome.

## Materials and methods

### Antibodies

The following antibodies were used in this study: rabbit anti-alpha tubulin 1:1000 for WB, rabbit anti-AP4E1 1:1000 for WB (a gift from Margaret Robinson, University of Cambridge; Hirst et al., 1999), rabbit anti-ATG9A 1:1000 for WB and 1:100 for IF (Abcam Cat# ab108338, RRID:AB_10863880), mouse anti-beta tubulin III (TUJ1) 1:800 for IF (Sigma-Aldrich Cat# T8660, RRID:AB_477590), rabbit anti-DAGLB 1:1000 for WB and 1:800 for IF (Abcam Cat# ab191159), chicken anti-GFP 1:500 for IF (Abcam Cat# ab13970, RRID:AB_300798), and sheep anti-TGN46 1:200 for IF (Bio-Rad Cat# AHP500, RRID:AB_324049). Horseradish peroxidase (HRP)-conjugated secondary antibodies were purchased from Sigma-Aldrich and used at 1:5000. Fluorescently labelled secondary antibodies used in this study were Alexa Fluor Plus 488-labelled goat anti-chicken IgY (Cat# A32931, RRID:AB_2762843), Alexa Fluor 488-labelled goat anti-mouse IgG (Cat# A-11029, RRID:AB_2534088), Alexa Fluor 568-labelled donkey anti-rabbit IgG (Cat# A10042, RRID:AB_2534017), and Alexa Fluor 680-labelled donkey anti-sheep IgG (Cat# A-21102, RRID:AB_2535755), all purchased from Thermo Fisher Scientific and used at 1:500.

### Cell culture

HeLa M cells were from ECACC. Other cell lines used in this study were previously described and are as follows: *AP4B1* knockout HeLa (clone x2A3); wild-type AP4B1-rescued *AP4B1* knockout HeLa (both described in Frazier et al., 2016); AP4B1-depleted SH-SY5Y, AP4E1-depleted SH-SY5Y, and their Cas9-expressing parental cells (mixed populations); wild-type HeLa and *AP4B1* knockout HeLa stably expressing RUSC2-GFP (mixed populations); endogenously-tagged HeLa SERINC1-Clover (clone A3) and HeLa SERINC3-Clover (clone B6) (all described in Davies et al., 2018).

HeLa and SH-SY5Y cells were maintained in Dulbecco’s Modified Eagle’s Medium (DMEM) high glucose supplemented with 10% v/v foetal calf serum, 4 mM l-glutamine, 100 U mL^-1^ penicillin and 100 μg mL^-1^ streptomycin and all cells were cultured at 37 °C under 5% CO_2_. Differentiation of SH-SY5Y cells into a neuronlike state was by sequential culture in the presence of retinoic acid and BDNF (Encinas et al., 2000). Cells were seeded at 25% confluency in standard culture medium and the following day the medium was replaced with medium containing 10 μM all-*trans*-Retinoic acid (Sigma-Aldrich Cat# R2625; 10 mM stock in DMSO). Two days later cells were split 1:4 and cultured for 3 further days in the presence of retinoic acid (5 days total). Cells were then washed in PBS and cultured for 5 days in serum-free DMEM containing 50 ng/ml BDNF (Miltenyi Biotec Cat# 130-096-285; 50 μg/ml stock in sterile ddH_2_O).

No cell lines used in this study were found in the database of commonly misidentified cell lines that is maintained by ICLAC and NCBI Biosample. The cell lines were routinely tested for the presence of mycoplasma contamination using DAPI to stain DNA and a PCR-based mycoplasma test.

### iPSC-derived cortical neurons

The protocol for generating human iPSCs was approved by the Institutional Review Board at Boston Children’s Hospital (IRB#: P00016119). iPSCs from an individual with *AP4B1*-associated SPG47 (*AP4B1*, NM_001253852.1: c.1345A>T / c.1160_1161delCA) and their sex-matched parent (heterozygous carrier, clinically unaffected; *AP4B1*, NM_001253852.1: c.1160_1161delCA / wild-type) were generated, maintained and differentiated into neurons as described previously (Behne et al., 2020; Teinert et al., 2019). iPSCs were maintained in StemFlex medium (Thermo Fisher Scientific, Cat# A3349401) on Geltrex (Thermo Fisher Scientific, Cat# A1413202), and passaged weekly with Gentle Cell Dissociation Reagent (STEMCELL Technologies, Cat# 07174). Cortical neurons were differentiated using induced NGN2 expression according to protocols modified from Zhang et al. (2013). Briefly, human iPSCs were dissociated into single cells with Accutase (Innovative Cell Technology, Cat# AT104–500) and seeded onto Geltrex-coated plates. On differentiation day −1, iPSCs were infected with concentrated rtTA-, EGFP-, and NGN2-expressing lentiviruses in presence of polybrene (Sigma-Aldrich, Cat# TR-1003-G). The next day, NGN2 expression was induced using doxycycline (Millipore, Cat# 324385), and infected cells were then selected using puromycin (Invitrogen, Cat# ant-pr-1) for up to 48 h. Growth factors BDNF (10 ng/ml, Peprotech Cat# 450–02) and NT3 (10 ng/ml, Peprotech, Cat# 450–03), and laminin (0.2 mg/L, Thermo Fisher Scientific Cat# 23017–015) were added in N2 medium for the first 2 days. Cells were then fed with BDNF (10 ng/ml), NT3 (10 ng/ml), laminin (0.2 mg/L), doxycycline (2 μg/ml), and Ara-C (2 μM, Sigma-Aldrich Cat# C1768) in B27 media and fed every other day until differentiation day 6. Cells were then dissociated with papain (Worthington, Cat# LK003178) supplemented with DNase (Worthington, Cat# LK003172) and replated with human iPSC-derived astrocytes (Astro.4U; Ncardia).

### Transfections and siRNA-mediated knockdown

Transient DNA transfections of wild-type *AP4B1* were carried out using a TransIT-HeLaMONSTER^®^ kit (Mirus Bio LLC), according to the manufacturer’s instructions, and a previously described construct, pLXIN_AP4B1 (Frazier et al., 2016).

Knockdown of AP-4 was achieved by combined siRNA targeting of *AP4E1* and *AP4M1* using ON-TARGETplus SMARTpools (*AP4E1*, L-021474-00; *AP4M1*, L-011918-01; Dharmacon), using a double-hit 96 hours protocol (Borner et al., 2012). For the first hit the final concentration of siRNA was 40 nM (20 nM *AP4M1* + 20 nM *AP4E1*). The second hit was performed 48 hours after the first hit with half the final concentration of siRNA. Transfections of siRNA were carried out with Oligofectamine (Thermo Fisher Scientific), according to the manufacturer’s instructions and control cells were transfected with ON-TARGETplus Non-targeting siRNA #1 (D-001810-01, Dharmacon).

### Western blotting

Estimations of protein concentrations were made using a Pierce BCA Protein Assay Kit (Thermo Fisher Scientific Cat# 23225). Cells were lysed for Western blot analysis in 2.5% (w/v) SDS/50 mM Tris pH 8 and incubated at 65 °C for 3 min. Lysates were then sonicated in a Bioruptor (fifteen 30 seconds on/off cycles at maximum intensity) to ensure complete solubilisation and fragmentation of nucleic acids. Samples were heated at 72 °C for 10 min in NuPAGE LDS Sample Buffer (Thermo Fisher Scientific Cat# NP0007). Samples were loaded at equal protein amounts for SDS-PAGE, performed on NuPAGE 4–12% Bis–Tris gels in NuPAGE MOPS SDS Running Buffer (all Thermo Fisher Scientific). PageRuler Plus Prestained Protein Ladder (Thermo Fisher Scientific) was used to estimate the molecular size of bands. Proteins were transferred to nitrocellulose membrane by wet transfer and membranes were blocked in 5% w/v milk in PBS with 0.1% v/v Tween-20 (PBS-T). Primary antibodies (diluted in 5% BSA in PBS-T) were added for at least 1 h at room temperature, followed by washing in PBS-T, incubation in secondary antibody (in 5% milk) for 30 min at room temperature, washing in PBS-T and finally PBS. Chemiluminescence detection of HRP-conjugated secondary antibody was carried out using Amersham ECL Prime Western Blotting Detection Reagent (GE Healthcare). Where representative blots are shown, the experiment was repeated at least two times.

### Immunofluorescence

For widefield microscopy, cells were grown onto 13 mm glass coverslips. For super resolution structured illumination microscopy (SR-SIM), cells were grown onto 18 mm precision glass coverslips (Marienfeld Cat# 0117580) that had been precleaned in 1 M HCl for 30 mins, washed twice in ddH2O and sterilised in 100% v/v ethanol. Cells were fixed in 3% formaldehyde in PBS (137 mM NaCl, 2.7 mM KCl, 10 mM Na_2_HPO_4_ and 1.76 mM KH_2_PO_4_, pH 7.4), permeabilised with 0.1% saponin in PBS and blocked in 1% BSA/0.01% saponin in PBS. Primary antibody (diluted in BSA block) was added for 1 hour at room temperature. Coverslips were washed three times in BSA block and then fluorochrome-conjugated secondary antibody was added in block for 30 min at room temperature. Coverslips were then washed three times in PBS and stained with 300 nM DAPI in PBS for 5 min, before a wash in PBS and a final wash in ddH2O. Apart from iPSC-derived cortical neurons, coverslips were mounted in SlowFade™ Glass Soft-set Antifade Mountant (Thermo Fisher Scientific Cat# S36917) and sealed with CoverGrip™ Coverslip Sealant (Biotium Cat# 23005). For iPSC-derived cortical neurons, coverslips were mounted in ProLong™ Diamond Antifade Mountant (Thermo Fisher Scientific Cat# P36965) and sealed with clear nail polish. For SR-SIM, glass slides were precleaned as described above for the glass coverslips. Channel alignment slides were prepared alongside the SR-SIM samples using multi-coloured fluorescent TetraSpeck™ 0.1 μm Microspheres (Thermo Fisher Scientific Cat# T7279).

### Widefield imaging

Widefield images were captured on a Leica DMi8 inverted microscope equipped with an iTK LMT200 motorised stage, a 63x/1.47 oil objective (HC PL APO 63x/1.47 OIL) and a Leica DFC9000 GTC Camera, and controlled with LAS X (Leica Application Software X). For DAGLB in RUSC2 overexpressing cells, images are representative of two technical replicates of separate coverslips from the same batch of cells, including three independent rescue transfections, with immunofluorescence and microscopy performed independently for each replicate. At least ten images were captured per condition, with both cell selection and manual focus performed on the RUSC2-GFP channel, without viewing the DAGLB channel. Consistent results were observed for cells expressing RUSC2 with an N-terminal GFP tag and with *AP4E1* knockout. For quantification of DAGLB at the TGN in HeLa and SH-SY5Y (undifferentiated and differentiated), experiments were performed in biological triplicate, from separate dishes of cells, and with immunofluorescence and microscopy performed independently for each replicate. Cells were selected for imaging using the DAPI channel only in the Navigator software module of LAS X. For HeLa, 30 images were captured per cell line per replicate, using autofocus on the DAGLB channel. For undifferentiated SH-SY5Y, 20 images were captured per cell line per replicate, with manual focus on the DAGLB channel. For differentiated SH-SY5Y, a minimum of ten images were captured per cell line per replicate, with manual focus on the DAGLB channel. For quantification of DAGLB at the TGN in iPSC-derived cortical neurons, the experiment was performed in technical triplicate with three separate coverslips per cell line from the same batch of cells and with microscopy conducted in three separate sessions. Neurons were selected for imaging using the TUJ1 signal to distinguish neuron cell bodies from astrocytes, without viewing the DAGLB channel. A minimum of 24 images were captured per cell line per replicate, with manual focus on the TGN46 channel.

### Structured illumination microscopy

Super resolution structured illumination Microscopy (SR-SIM) was performed on a Zeiss Elyra PS.1 microscope equipped with a 63x/1.46 oil objective (alpha Plan-Apochromat 63x/1.46 Oil Korr M27) and a PCO pco.edge 4.2 sCMOS Camera, and controlled with Zeiss ZEN 2 software with the SR-SIM module. Imaging was performed sequentially, with SERINC1 or SERINC3 (labelled with Alexa Fluor Plus 488) imaged first and DAGLB or ATG9A (labelled with Alexa Fluor 568) imaged second, using 3 rotations of the grid pattern and 5 phases for each rotation. Experiments were performed in biological duplicate, from separate dishes of cells, and with immunofluorescence and microscopy performed independently for each replicate. 20 cells were imaged per condition per replicate, and cell selection and manual focus were performed on the first acquired channel only, without viewing the second channel. For channel alignment, slides with multi-coloured fluorescent beads were imaged before and after each experiment using the same acquisition settings. The data were processed using the SIM module in Zen 2 software in manual mode, using default settings and a theoretical PSF model, except for the following settings: Noise filter −1; SR frequency weighting 2; Max.Isotrop on; Baseline shifted on; Raw scale on. Channel alignment fit parameters were calculated in Zen 2 software using a multi-coloured bead image as input, in affine mode. The resulting parameters were tested on further bead images and then applied to the experimental images. The channel-aligned images were used for colocalisation analysis as described below.

### Colocalisation analysis

Colocalisation analysis was performed by comparing the fluorescent spots of the two proteins from super resolution SIM images, following processing and channel alignment as described above. Colocalisation of proteins was quantified in ImageJ (Schneider et al., 2012) using the Coloc2 plugin, included in the FIJI package distribution (Schindelin et al., 2012). Individual cells were cropped and manually outlined in FIJI using the SERINC1 or SERINC3 channel. After this step, the complete analysis pipeline was performed by means of a macro script, provided as Supplemental Material. The functions used at each processing step to identify the spots in the two channels are reported in the following, as described in ImageJ, with parameters in pixels. The pixel size of the images is 32 nm.

Each channel was pre-processed separately by first reducing the image noise (Gaussian Blur, sigma: 1) and then removing the background signal (Subtract Background, rolling ball radius: 20). The spots were identified as regions with intensity twice the standard deviation above the local average signal (Remove Outliers, block radius:40, standard deviations:2) and used to generate an initial mask. Small spots were discarded (Analyze Particles, size larger than 4) and the edges of the remaining spots were smoothed (Gaussian Blur, sigma:0.75). The resulting soft mask was applied to the original intensity image and finally the resulting images from each channel were provided as input to Coloc2. Colocalisation output metrics were Pearson’s Correlation Coefficient and Mander’s overlap coefficients M1 and M2.

### Quantification of DAGLB at the *trans*-Golgi network

The accumulation of DAGLB at the TGN was quantified by comparing the fluorescence average intensity in that region with the intensity in the rest of the cell, from widefield images. The analysis was performed with a workflow implemented in the software CellProfiler (Mcquin et al., 2018), version 3.1.9, slightly adapted to each cell type analysed. The pipeline files for each cell type are available as Supplemental Material.

The position and extension of each cell was determined by first identifying the nuclei, using the minimum cross entropy thresholding method on the DAPI channel. The area of each cell was then determined from the DAGLB channel, previously filtered to achieve a uniform distribution of intensity over the cell (two steps of the Smooth operation, first by average filtering, followed by median filtering). Using the nuclei detected as seeds, the area of the cells was obtained from the pre-processed image by watershed combined with Otsu thresholding. The area occupied in each cell by the TGN was determined from the TGN46 channel, after background subtraction, by minimum cross entropy thresholding. This information was used to determine the two complementary regions of each cell where to measure the average fluorescence intensity of the DAGLB signal, after illumination correction. From these measurements a ratio of DAGLB intensity between the TGN and the rest of the cell was calculated. For HeLa and iPSC-derived neurons, ratio values were calculated for each individual cell. For SH-SY5Y, ratio values were not calculated for individual cells, because they were too overlapping. Instead ratios were calculated for each image, based on the total cell area and total area occupied by the TGN across the entire field of view. In all cases, edge cells were excluded from the analysis.

The workflow was partially modified for the analysis of iPSC-derived cortical neurons, in order to exploit the additional fluorescence signal from the neuron cell body marker TUJ1. Specifically, the marker was used to discriminate neurons from astrocytes, also present in the field of view, and to determine the area of the cells of interest.

### Statistical analysis of microscopy data

Statistical analyses were performed in GraphPad Prism version 8.4.1 for Windows. Ratio data were log2 transformed. Data were analysed using a Kruskal-Wallis test with Dunn’s multiple comparisons test for comparisons of more than two conditions or a two-tailed Mann–Whitney *U*-test when only two conditions were compared. Replicate data were combined for plotting as there were no significant differences between replicates of the same condition.

### Sensitive statistical analysis of AP-4 knockout Dynamic Organellar Maps

We previously generated six organellar maps from HeLa (duplicates of wild-type, *AP4B1* knockout, and *AP4E1* knockout; Davies et al., 2018), and analysed them with our established MR (movement and reproducibility) test (Itzhak et al., 2019) to identify proteins with altered subcellular localisation. Briefly, for each protein, the abundance profiles from wild-type and knockout maps were subtracted pairwise, to obtain four sets of ‘delta profiles’ (wild-type-AP4B1, wild-type-AP4E1, both in duplicate). Proteins that did not shift localisation had ‘delta profiles’ close to zero. Proteins with significant outlier delta profiles in all replicates (high M(ovement) score, calculated as a robust Mahalanobis Distance within Perseus (Tyanova & Cox, 2018)) were candidate shifting proteins. As a further filter, the similarity of the direction of the shift across replicates was evaluated by calculating delta profile correlation (R(eproducibility) score). To define useful cut offs for M and R, the same analysis was performed on six wild-type HeLa maps (Itzhak et al., 2016). Here, no genuine shifts were expected, so the number of hits identified at a given set of MR cut offs allowed an estimate of the false discovery rate (FDR) for hits detected in the AP-4 knockout vs wild-type analysis. In Davies et al. (2018), we used extremely stringent MR settings to filter the data: from the four M scores and the two R scores, we chose the respective lowest values (i.e. most conservative estimate of protein movement). The estimated FDR was <1%, and indeed, all three identified hits were subsequently validated by imaging (Davies et al., 2018). Here, we reanalysed this dataset with less stringent filters. We calculated the median M score from the four replicates, and the average of the two R scores. The same calculations were then performed for the six wildtype HeLa maps to estimate the FDR. We chose the same M-score cut-off as previously (M>4), and an R-score cut-off of >0.92 (the highest value that still included all three previously identified hits). As a further filter, we required the delta profiles of hits to have a correlation >0.9 across both AP-4 knockout clones; this filter was not applied to the wild-type-only dataset. In addition to the previously identified three proteins, we thus identified eight new hits, with an estimated FDR of 25%.

### Statistical analysis of TEPSIN-GFP immunoprecipitations

Sensitive TESPIN-GFP immunoprecipitation SILAC MS data were acquired in our previous study (Davies et al., 2018). The processed and filtered dataset described was reanalysed in Perseus version 1.6 (Tyanova et al., 2016). A one-sample *t*-test (two-tailed) was applied to compare the *L/H* ratios for each protein to zero (null hypothesis of no change between TEPSIN-GFP immunoprecipitations and mock immunoprecipitations from wild-type HeLa cells).

### Data availability

All data supporting this work are available on reasonable request to the corresponding author.

## Acknowledgements

We thank Margaret (Scottie) Robinson for her generous sharing of reagents and resources, support, and helpful discussions. We thank Matthias Mann for his continued support of this project, Jennifer Hirst for sharing of reagents, and Julia Schessner and members of the Robinson and Mann Labs for valuable feedback. Special thanks to the MPIB Imaging Facility for outstanding technical support, in particular to Giovanni Cardone for his advice and assistance with implementation of the image analysis pipelines used in this study and to Martin Spitaler for his expert technical advice for imaging experiments. This work was funded by the German Research Foundation (DFG/Gottfried Wilhelm Leibniz Prize MA 1764/2-1), the Max Planck Society for the Advancement of Science, and a Humboldt Research Fellowship to A.K.D. The authors declare no competing financial interests.

## Author Contributions

G.H.H.B. and A.K.D. conceptualised and designed the experiments. A.K.D. performed and analysed most of the experiments. G.H.H.B. performed statistical analysis of the Dynamic Organellar Maps data. D.E.-F., M.Z., H.J. and W.A.S. performed the iPSC experiments. A.K.D. and G.H.H.B. wrote the original and revised manuscripts, with contributions from the other authors. G.H.H.B. supervised the project.

## Supplementary Figure Legends

**Figure S1.**
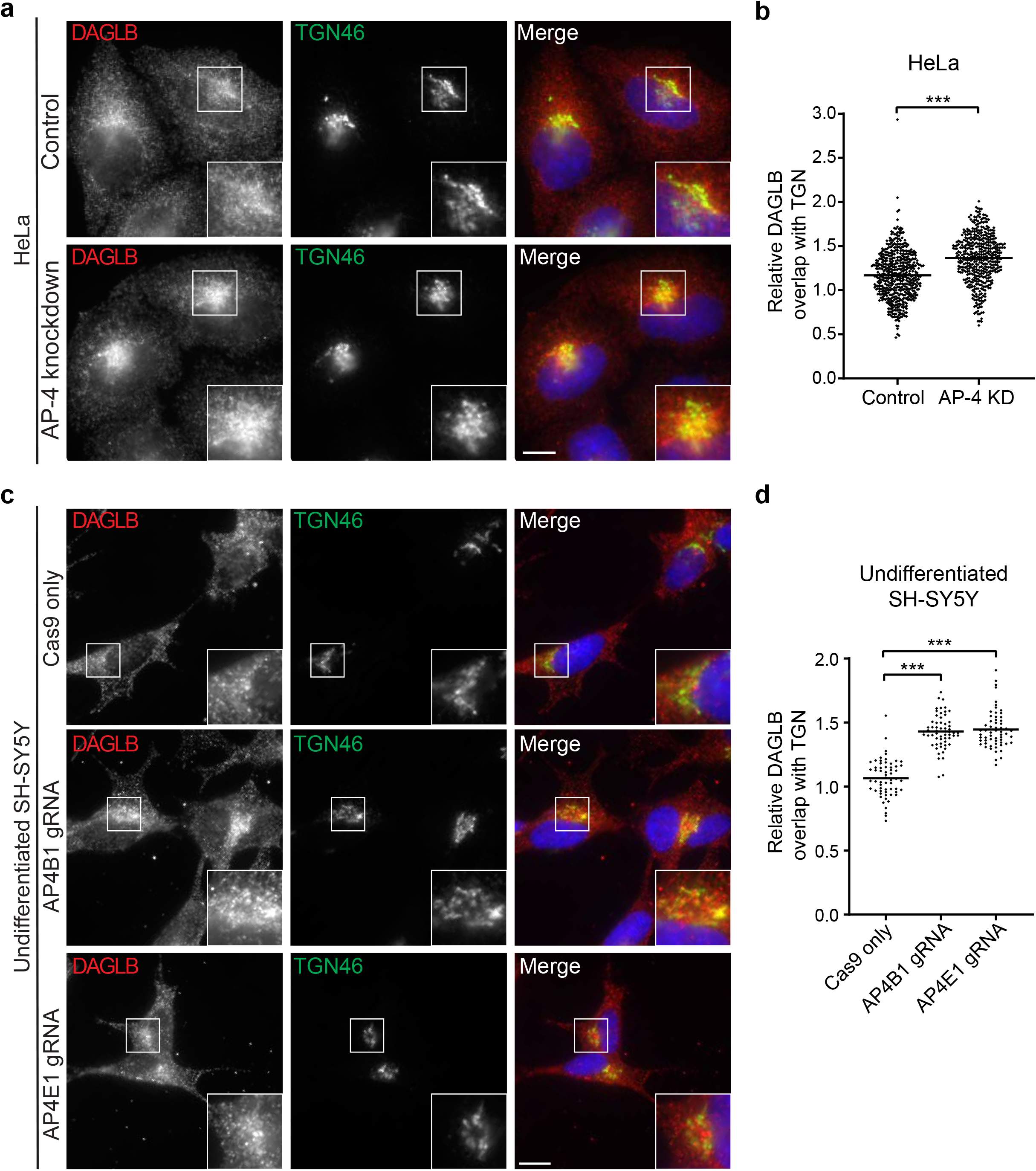
DAGLB accumulates at the *trans*-Golgi network (TGN) in AP-4 knockdown HeLa and undifferentiated AP-4 depleted SH-SY5Y cells. **(a)** Widefield imaging of immunofluorescence double labelling of DAGLB and TGN46 in HeLa cells transfected with a non-targeting siRNA (control) or with siRNA to knock down AP-4. In the merged image, DAPI labelling of the nucleus is also shown (blue). Scale bar: 10 μm. **(b)** Quantification of the ratio of DAGLB labelling intensity between the TGN and the rest of the cell, in the cells shown in **a**. The experiment was performed in biological triplicate, and >140 cells were scored per cell line in each replicate. The graph shows combined replicate data, and each datapoint indicates the ratio for an individual cell (mean indicated). Data were subjected to a two-tailed Mann–Whitney *U*-test: \*\*\**p* ≤ 0.001. **(c)** Widefield imaging of immunofluorescence double labelling of DAGLB and TGN46 in control (parental Cas9-expressing), AP4B1-depleted and AP4E1-depleted undifferentiated SH-SY5Y cells. In the merged image, DAPI labelling of the nucleus is also shown (blue). Scale bar: 10 μm. **(d)** Quantification of the ratio of DAGLB labelling intensity between the TGN and the rest of the cell, in the cells shown in **c**. The experiment was performed in biological triplicate, and 20 images were quantified per cell line in each replicate. The graph shows combined replicate data, and each datapoint indicates the ratio calculated from a single image (mean indicated). Data were subjected to a Kruskal-Wallis test with Dunn’s Multiple Comparison Test for significance: \*\*\**p* ≤ 0.001.

**Table S1.** Summary of AP-4 vesicle cargo and accessory proteins supported by unbiased proteomic approaches. Inclusion criteria: evidence from at least two orthogonal experiments, where at least one was a hypothesis-free proteomic approach assaying AP-4-association with endogenous proteins. Candidates requiring validation are in grey.

| Gene name | Role | Proteomic evidence | Other evidence | References |
| --- | --- | --- | --- | --- |
| **ATG9A** | Cargo | Maps, IP, BioID, vesicle fraction | Microscopy, *in vitro* interaction | Davies et al. 2018; Ivankovic et al. 2020; Mattera et al. 2015/2017 |
| **SERINC1** | Cargo | Maps, IP, vesicle fraction | Microscopy | Davies et al. 2018 |
| **SERINC3** | Cargo | Maps, IP, vesicle fraction | Microscopy | Davies et al. 2018 |
| **DAGLB** | Cargo | Maps, IP, vesicle fraction | Microscopy | This study; Davies et al. 2018 |
| **TEPSIN** | Accessory (unknown function) | IP, BioID, vesicle fraction | Microscopy, *in vitro* interaction | Borner et al. 2012; Davies et al. 2018; Frazier et al. 2016; Mattera et al. 2015 |
| **RUSC1** | Accessory (anterograde transport adaptor) | Vesicle fraction, membrane fraction |  | Davies et al. 2018 |
| **RUSC2** | Accessory (anterograde transport adaptor) | BioID, vesicle fraction | Microscopy, *in vitro* interaction | Davies et al. 2018 |
| **HOOK1** | Accessory (retrograde transport adaptor, FHF complex) | IP, BioID | *In vitro* interaction | Davies et al. 2018; Mattera et al. 2020 |
| **HOOK2** | Accessory (retrograde transport adaptor, FHF complex) | None, but inferred from HOOK1/3 | Microscopy, *in vitro* interaction | Mattera et al. 2020 |
| **HOOK3** | Accessory (retrograde transport adaptor, FHF complex) | IP, BioID | *In vitro* interaction | Davies et al. 2018; Mattera et al. 2020 |
| **FAM160A1** | Accessory (retrograde transport adaptor, FHF complex) | IP, BioID | Microscopy | Davies et al. 2018; Mattera et al. 2020 |
| **FAM160A2** | Accessory (retrograde transport adaptor, FHF complex) | IP, BioID | Microscopy, *in vitro* interaction | Davies et al. 2018; Mattera et al. 2015/2020 |
| **AKTIP** | Accessory (retrograde transport adaptor, FHF complex) | IP, BioID |  | Davies et al. 2018; Mattera et al. 2020 |
| **AAGAB** | Accessory (unknown function, also binds AP-1 and AP-2) | IP, BioID |  | Davies et al. 2018 |
| **PTPN9** | Candidate accessory, needs further validation | Maps |  | This study |
| **LNPEP** | Candidate cargo, needs further validation | Maps |  | This study |

Maps: protein had a significantly altered distribution in AP-4 knockout cells in 'Dynamic Organellar Maps' fractionation-based spatial proteomics; IP: protein co-immunoprecipitated with the AP-4 complex; BioID: protein was specifically biotinylated by AP-4 complex baits; vesicle fractionation: protein had an altered abundance in vesicle fractions prepared from AP-4-deficient cells. Integral membrane proteins of AP-4 vesicles are considered cargo; accessory factors are cytosolic proteins that transiently associate with AP-4 vesicles. The tetrameric AP-4 complex itself (AP4E1, AP4B1, AP4M1, AP4S1) is not listed here.

